# Human *in vivo* midtarsal and subtalar joint kinematics during walking, running, and hopping

**DOI:** 10.1101/2023.09.15.558017

**Authors:** Anja-Verena Behling, Lauren Welte, Luke Kelly, Michael J Rainbow

## Abstract

The interaction among joints of the midtarsal complex and subtalar joint is essential role for locomotor function; however, its complexity poses substantial challenges in quantifying their motions. We determine the mobility of these joints across locomotion tasks and investigate their alignment with individual talus morphology.

Utilizing highly accurate biplanar videoradiography, three-dimensional bone kinematics were captured during walking, running, and hopping. We calculated the axis of rotation of each midtarsal and subtalar joint for the landing and push-off phases, respectively. A comparison was made between these rotation axes and the morphological subtalar axis. Measurement included total rotation about, the orientation of the rotation axes in the direction of the subtalar joint and its deviation via spatial angles for both phases.

The rotation axes of all three bones relative to the talus closely align with the morphological subtalar axis. This suggests that the midtarsal and subtalar joints’ motions might be described by one commonly oriented axis. Despite having such axis, the location of axes and ranges of motion differed among the bones.

Our results provide a novel perspective of healthy foot function across different sagittal plane-dominant locomotion tasks underscoring the importance of midtarsal and subtalar motion with respect to subject-specific talus morphology.

## 1. Introduction

The movement of the midtarsal joint complex (talonavicular and calcaneocuboid joints) together with the subtalar joint plays an important role in facilitating bipedal gait [1–5]. However, quantifying *in vivo* midtarsal and subtalar joint motion is challenging. The rare data that has been captured has often been used to test the incorrect assumption that the foot becomes rigid during push-off [6–9], which is still a common misconception of foot function. It was previously suggested that the large variation in bone shape has the potential to modify midtarsal complex and subtalar joint function [10–13].We propose to investigate the function of the midtarsal complex and subtalar joint under a more modern framework that views the foot as a highly mobile structure [6], while accounting for the underlying morphology. This approach may provide new insights into foot function.

Different approaches have been established to understand the midtarsal complex and subtalar joint. Publications based on high-resolution data from intracortical bone-pins [14–16] and video radiography [7,17,18] provide unique insights into how the midtarsal and subtalar joints are moving throughout walking and slow running. These studies highlight the complex interactions between these bones and display large inter-subject variation, questioning previous conceptions of the foot as a rigid unit. Intracortical bone-pin studies typically rely on a static reference pose, and kinematics were calculated using either Euler angles [16,19] or helical axis of motion (rotation axis) approaches [14,15] from optical motion capture data. While these approaches allowed for the calculation of bone kinematics relative to a reference bone, they did not address large inter-individual variation in morphology. This is important as it allows us to include variations in morphology to explain differences in foot function. Moreover, whether there is a common pattern in terms of axis orientation has never been investigated *in vivo* [20], despite the utilization of helical axis approach for some of the bone-pin studies [14,15] and a comparison of rotation axes would have been feasible. However, the highly invasive nature of intracortical bone-pins makes it impossible to establish this approach as a common technique and non-invasive alternatives are required.

Dynamic imaging with biplanar video radiography (BVR) directly captures bone morphology in motion without invasive hardware, providing insight into the relationship between morphology and function. A recent single-plane video radiography study reported by Wang and colleagues [17] described the motion of the talonavicular, calcaneocuboid, and subtalar joints as synchronous and homodromous in all three planes during overground walking. This might suggest that these joints share a common rotation axis about which they move, as previously suggested by Elftman and Manter [21]. To address the question of a common rotation axis, we employed the helical axis of motion approach, defining a rotation axis based on the motion between two bones and two time points. Understanding the motion of the midtarsal complex and subtalar joint in terms of a common rotation axis using BVR is useful for two reasons: first, it allows us to compare bone motions directly, relative to a single reference bone’s coordinate system, providing us with the opportunity to directly compare the rotation axes across bones, participants, and locomotion tasks. Second, rotation axes can be tied to the underlying bone morphology.

The anatomical features of articular surfaces can be used to calculate a morphological axis. A morphological axis defined by the calcaneus and navicular facets on the talus is likely a good candidate for capturing the rotation axes of the bones in the midtarsal complex and subtalar joint as both articular facets directly engage with the calcaneus and navicular bones [21 p. 416ff, 22]. We define this axis as the morphological subtalar axis because it is thought to align with the subtalar joint rotation axis [21 p. 416ff, 22]. Conconi and colleagues [23] have previously compared morphology-based bone specific coordinate systems with mean helical axes of motion between pre-defined foot positions (achieved by wedges during upright standing). They showed that the dorsi-plantar flexion axis in the talocrural joint coincides well with a cylinder-based axis through the talar dome (mean angle difference of 12.3°) and their subtalar joint axis defined by calcaneal posterior and middle talar facets are within a 17.2° agreement with the mean rotation axis of the subtalar joint. Thus, if the midtarsal complex and subtalar joint likely rotate around a common axis [21,24], then we would expect this axis to align with the morphological subtalar axis [23]. This analysis deviates from previous work that examines talonavicular and calcaneocuboid motion independently [5,14,16,25]. Our rationale here is that it has been shown previously that the navicular and cuboid move as a unit [7,15– 17,25]. Analyzing their motion relative to the talus allows us to directly test whether this is the case.

Understanding how the motion of the midtarsal and subtalar joints relate to each other and to talus morphology, and how its overall function relates to morphology may allow researchers to determine which shape features are important to healthy human gait. This may provide insight into evolutionary origins. Moreover, recent statistical shape modeling studies [28–32] highlighted substantial morphological variations of non-pathological human tali. These shape variations have considerable influence on the talonavicular and subtalar joint facets [11,13,26]. Hence, if morphological features in these facets change, so might the morphological axis and therefore the rotation axes and resulting kinematics. Understanding these midtarsal complex and subtalar joint form-function relationships may allow scientists and clinicians to better understand the role of an individual’s anatomy, or changes to it, in the onset and progression of pathology. Moreover, we also aim to understand if the initially suggested synchronous and homodromous motion of the midtarsal complex and subtalar joint during walking [17] extends to other dynamic locomotion tasks (such as running and hopping) by quantifying the alignment of the rotation axes during these tasks. Running and hopping were selected for two reasons, both tasks are common during daily activities and part of many sports. Additionally, hopping is a locomotion task with large range of motion in the midtarsal complex and subtalar joint.

The purpose of this study was to determine if 1) midtarsal and subtalar motion occurs about a commonly oriented rotation axis in a range of locomotor tasks and 2) whether the individual midtarsal and subtalar joint axes align with the morphological subtalar axis. We utilize highly accurate BVR technology to obtain bone kinematics and calculate each bone’s motion, relative to the talus, using the finite helical axis of motion approach to represent the rotation axes [27,28].

## 2. Methodology

### 2.1. Overview and Participants

Kinematic data were combined from two separate data collections and study protocols for this study. All participants provided written informed consent (MECH-061-17 and MECH-063-18). There was no overlap in participants between the two data sets.

Walking and running data were obtained from nine healthy participants (5 females; mean ± std; mass 76.1 ± 15.8 kg; height 171.1 ± 15.8 cm; 20.0 ± 6.8 years) with no history of lower limb injury. Participants walked and ran at self-selected speeds while wearing minimalist shoes (Xero, Prio). BVR images were collected from the right leg of each participant (125 Hz for walking, 250 Hz for running, range of 70-80 kV, 100-125 mA). Ground reaction forces were collected simultaneously using two force plates (1125 Hz, AMTI Optima, AMTI, Watertown, MA), whose forces were combined and synchronized to the BVR system via a trigger step function. We analyzed one gait cycle per condition from three collected trials for each participant. The field of view for the BVR system is relatively small, so the foot bones of interest were not always visible for the entire stance phase. Therefore, each trial was screened for the maximal number of frames in which the bones were visible (∼85-100% of stance; Supplement Figure S1).

In addition to walking and running, we captured unilateral hopping. Hopping was chosen because it produces a large range of cyclical motion about the ankle and midtarsal and subtalar joints [29,30] and can be captured within the small field of view of the BVR system. For the hopping data, nine healthy subjects (5 females; mean ± std, mass 71.3 ± 14.5 kg; height 171.4 ± 10.7 cm; 27.6 ± 7.4 years) with no history of lower limb injury provided written informed consent to participate in this study. Participants hopped barefoot on their right leg at a frequency of 156 beats per minute, guided by a metronome. The BVR system (125 Hz, range of 70–80 kV, 100–125 mA) and force plate data were collected simultaneously (1125Hz, AMTI Optima, AMTI, Watertown, MA) and manually synchronized to the BVR system using optical motion capture (Qualysis Track Manager, Goeteborg, Sweden). Three hops with the entire foot in the field of view were kept for further processing.

### 2.2. Data Processing: Kinematics Derived from Biplanar Video Radiography

A computed tomography scan (CT, 120 kV, 60mA, model: Lightspeed 16, n = 11, Revolution HD, n = 1, General Electric Medical Systems, USA) was obtained from the right foot while participants were lying in a prone position (average resolution: 0.356 × 0.356 × 0.625 mm). Talus, navicular, calcaneus, and cuboid bones were segmented to create three-dimensional bone models (Mimics 24.0, Materialise, Leuven, Belgium). Subsequently, bone surface meshes and partial volumes were generated, and digitally reconstructed radiographs were created [31].

The processing pipeline for foot BVR data can be found in our earlier research [32,33]. Briefly, the highspeed cameras were calibrated using a custom calibration cube, and the images were undistorted in XMALab (Brown University, USA) [34,35]. The translation and orientation of the bones of interest were measured by tracking partial volumes representing each bone of interest using Autoscoper (Brown University, USA) [31].

Gait events were defined using a 15N threshold in the vertical ground reaction force to determine touchdown and toe-off during locomotion. As walking and running have distinct energetic functions for braking and propulsion phases [36], we have split the stance phase into two phases according to the zero-crossing of the anterior-posterior ground reaction force [36]. The zero-crossing point (transition point) does not exist in the hopping data set as the participants were asked to hop on the spot. Hence, the transition point for the hopping data set was defined as the maximal vertical ground reaction. To keep the terminology consistent across all locomotion tasks, the landing phase is defined from touchdown to the transition point and the push-off phase is defined from the transition point to toe-off.

We implemented a morphological coordinate system for the talus, which was inspired by previously published morphology-based coordinate systems [23,37]. Two spheres were fitted to the talus (Figure 1), one to the calcaneus and the other to the navicular facet. Our morphological subtalar axis (z-axis) orientation was defined by the vector that connected the centroids of each sphere (subtalar vector) and oriented anteriorly and superiorly [38]. We chose to define the subtalar axis in accordance with Modenese and colleagues’ approach [37] as it includes two morphological sites on the talus. Our approach differs from Conconi et al.’s method [23], who determined the morphological subtalar axis as with two spheres fitted to the calcaneus facets. The y-axis was the mutual perpendicular of the long axis of a cylinder fit to the talar dome and the z-axis and is oriented laterally for a right foot. Lastly, the x-axis was calculated as the cross product of the y- and z-axes. The origin lies on the cylinder axis where the distance to the subtalar vector line is minimal. Coordinate systems for the calcaneus, navicular and cuboid are not needed as rotation axes are resolved into one common coordinate system (the talus coordinate system).

**Figure 1:**
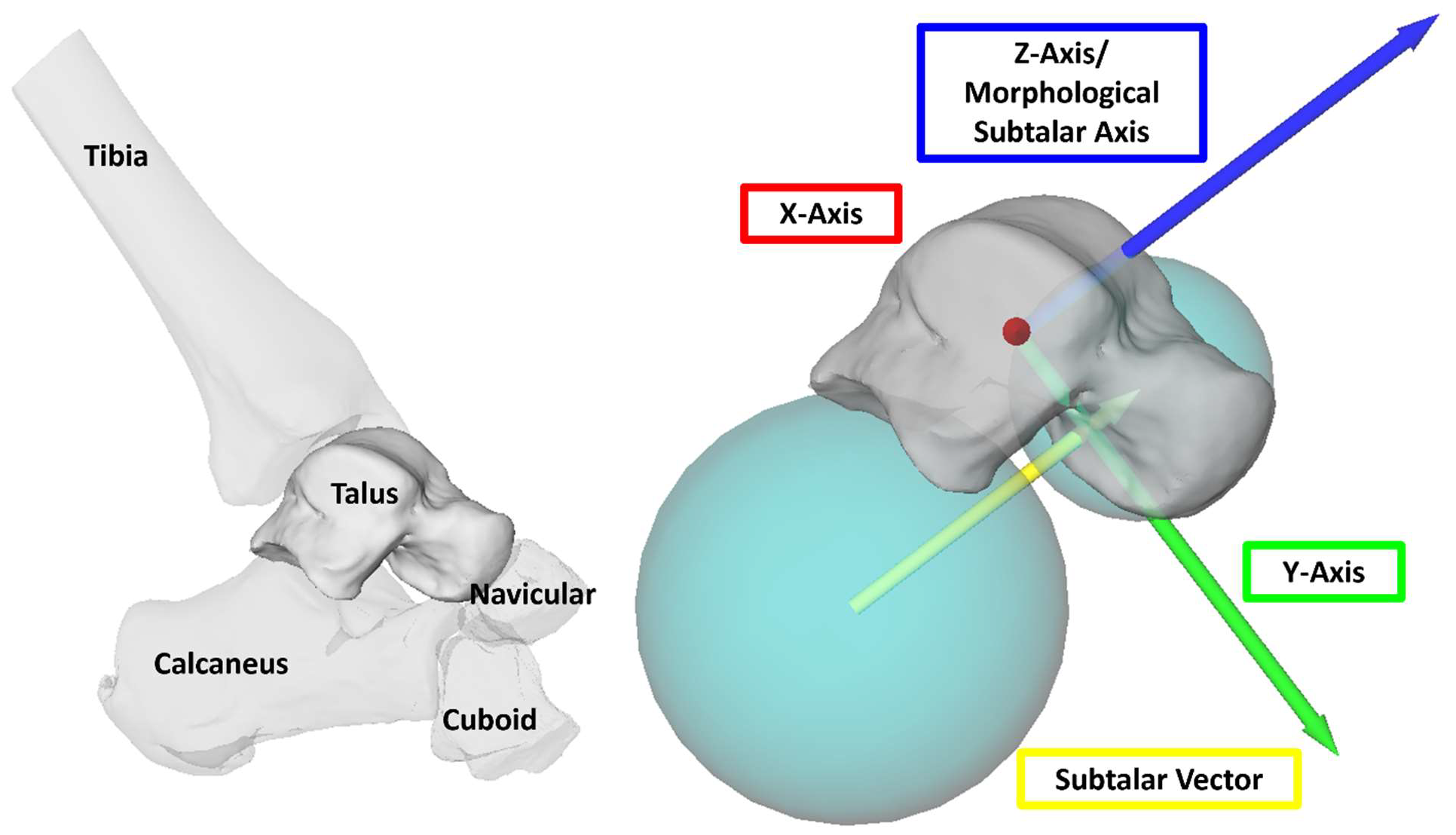
Right foot; lateral view. Left panel shows the talus and its surrounding bones. Right panel shows an exemplary talus coordinate system for one participant. X-axis is red (oriented out of the page, laterally), y-axis green (anteriorly-inferiorly oriented) and z-axis blue (morphological subtalar joint axis; oriented anteriorly-superiorly). The morphological subtalar axis is parallel to the vector connecting the centroids of the sphere fits of the talonavicular and subtalar facet (teal spheres). The origin is defined on the cylinder line and is located at the point closest to subtalar vector (yellow arrow).

Finite helical axes of motion were calculated for the bones of interest [27,28], and herein is termed rotation axis. The rotation axis represents the change in position and orientation of one rigid body moving relative to a mathematically fixed rigid body (pose) – here, the talus. It defines an axis in space that the moving body rotates about and translates along. The rotation axis might not be identical with the joint axis per instant in time (instantaneous helical axis). We focus on the orientation of the rotation axis in the z-direction and the angle swept by the moving body as it rotates about the rotation axis (total rotation).

We calculated the rotation axis orientation and total rotation of the subtalar joint (calcaneus relative to talus), talonavicular joint (navicular relative to talus), and talocuboid joint (cuboid relative to talus) and resolved the axis in the talus coordinate system (Figure 2). Each gait phase (landing and push-off) corresponds to one rotation axis, which was calculated as the difference in pose between the two time instants (touchdown and transition point for the landing phase and between the transition point and toe-off for the push-off phase). If a rotation axis reported a total rotation ≤5° or if the bone (typically the cuboid) was occluded by other bones during tracking, we excluded this helical axis of the bone for the participant from further analysis. If bones were not trackable at touchdown or toe-off, we used the next available frame to calculate the landing and push-off phase, respectively. We also calculated the spatial angle between the z-component of the morphological subtalar axis and each bone’s rotation axis to provide an alternative measure of alignment between the rotation axes and the morphological subtalar axis.

**Figure 2:**
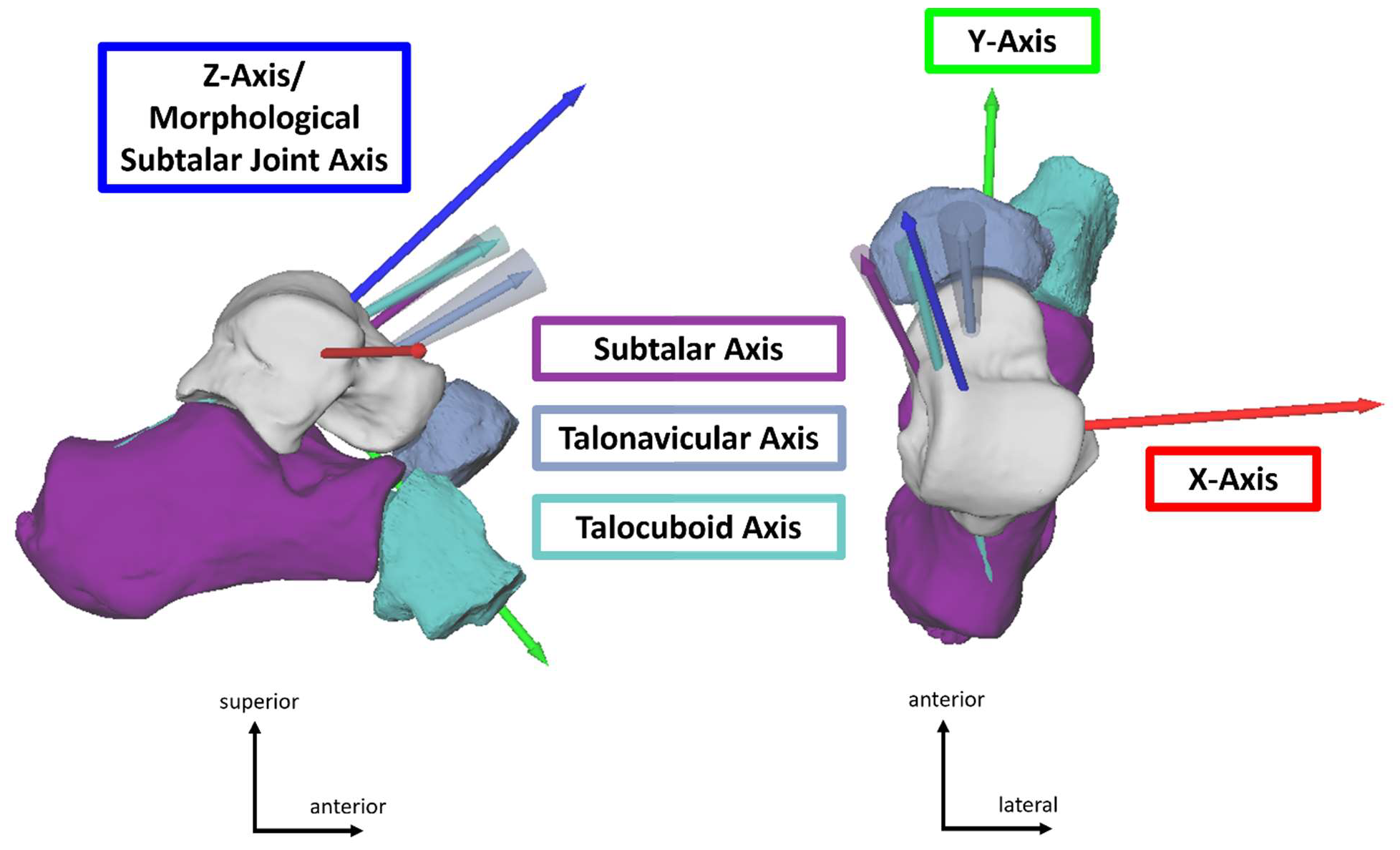
Right foot, lateral (left) and superior (right) view. Representative rotation axes for the push-off phase during hopping for all three joints (subtalar in purple, talonavicular in blue, and talocuboid in teal) relative to the talus (for n_Participants_ = 1). Rotation axes are indicated as arrows with their standard deviation displayed as cones about each axis. The talus coordinate system is indicated in red (x-axis), green (y-axis) and blue (z-axis/ morphological subtalar axis).

### 2.3. Analysis

In general, we use Bland-Altman plots to assess similarity. To explore systematic differences in the data set, we employ one-way ANOVA tests, with the caveat that due to our small sample size, the statistical power is small and should be viewed with caution. The following statistical analyses should be viewed with caution, as they are exploratory.

To determine whether the rotation axes are similarly oriented, we performed one-way ANOVAs as a proxy for similarity measures (absence of difference) for the z-axis orientation component of the rotation axes of the three joints relative to each other (Supplement Table S1-3). Additionally, we created Bland-Altman plots as a measure of agreement to test whether the three rotation axes agree in their orientation in the z-direction (Supplement Figure S2-4).

To determine whether the rotation axes were aligned with the morphological subtalar axis, we again examined the z-component of the rotation axes’ orientation, which corresponded to the morphological subtalar axis direction. If a rotation axis orientation has a value of 1 in the subtalar joint direction (z-direction), it means that the rotation axis is perfectly aligned with the morphological subtalar joint axis (same orientation), and all its motion occurs about this axis.

To examine whether the navicular and cuboid moved as a unit relative to the talus, and whether their motion was greater than subtalar motion as shown previously, we performed one-way ANOVAs comparing the magnitude of total rotations across the different bones.

Differences between landing and push-off phases regarding total rotation magnitudes and orientation in the *z-*axis per joint were determined using paired t-tests for each mode of locomotion and joint.

Non-parametric equivalents were used whenever assumptions of paired t-tests or ANOVAs were not met. The significance level was set to 0.05 and adjusted when required using a Bonferroni correction.

## 3. Results

Since there were no differences between landing and push-off phase in total rotation and orientation for any of the three bones for all three locomotion tasks (p ≥ 0.039, alpha = 0.0176), we only report push-off results. Please refer to the supplements for landing (Supplement Figures S2-7, Tables S1-3).

Despite large variation in ranges of motion across joints and participants (Figure 3), we found that all three rotation axes were largely aligned with each other and the morphological subtalar axis across locomotor modes (Figure 4 & 5, Table 1, Supplement Figures S2-4). Moreover, the spatial angles between the morphological subtalar axis and the rotation axes were between 13.0-25.8° (subtalar), 16.3-24.9° (talonavicular), and 18.7-23.5° (talocuboid) across both gait phases and all three locomotion tasks (Supplement Table S4).

**Table 1:** Mean (standard deviation) of total rotation and z-component of the rotation axes of the talonavicular (navicular), talocuboid (cuboid), and subtalar (calcaneus) joints during push-off.

|  | Total Rotation (°) |  |  | Morphological Subtalar Alignment |  |  |
| --- | --- | --- | --- | --- | --- | --- |
|  | Walking | Running | Hopping | Walking | Running | Hopping |
| **Navicular** | 19.5 (9.2) | 19.5 (5.0) | 15.4 (4.3) | 0.92 (0.11) | 0.86 (0.12) | 0.92 (0.04) |
| **Cuboid** | 17.1 (9.1) | 16.8 (1.0) | 13.7 (3.8) | 0.95 (0.03) | 0.91 (0.06) | 0.92 (0.05) |
| **Calcaneus** | 10.8 (4.0) | 10.4 (3.6) | 10.0 (2.2) | 0.90 (0.06) | 0.86 (0.12) | 0.95 (0.03) |

**Figure 3:**
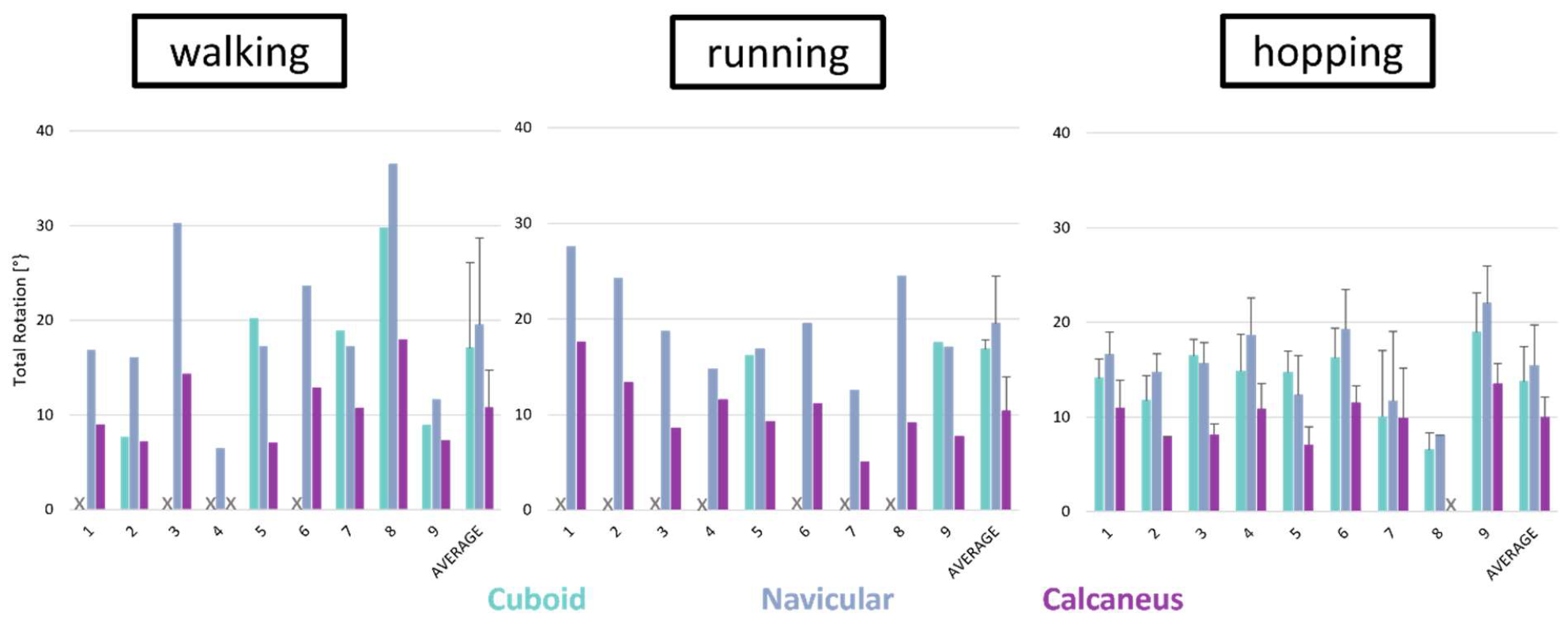
Total rotation in degrees during push-off phase of different modes of locomotion for the talocuboid (cuboid; teal), talonavicular (navicular; blue), and subtalar (calcaneus; purple) joints for all participants (n_Participants_ = 9; n_Trials_walk/run_ = 1; n_Trials_hopping_ = 3). Participants’ total rotation for some bones might be missing if the bone could not be tracked or was ≤ 5°. In these instances, there are no data presented in the graphs (indicated with an X).

**Figure 4:**
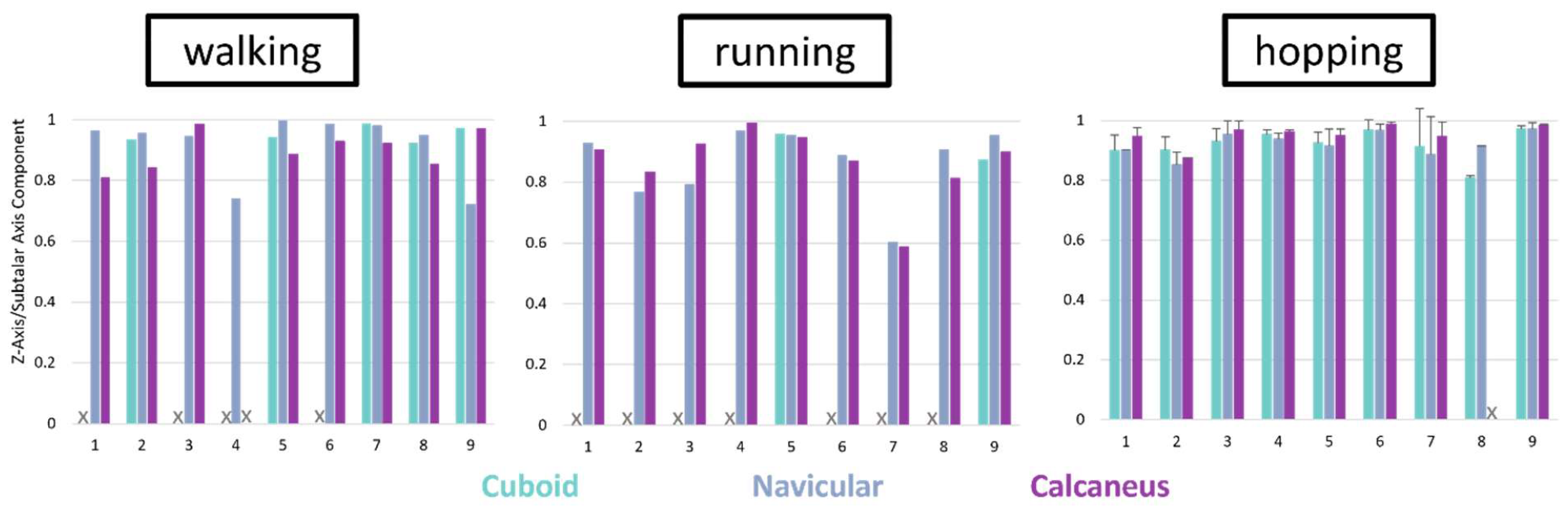
Orientation of rotation axes in the morphological subtalar axis direction (z-axis) during push-off phase of different modes of locomotion for the talocuboid (cuboid; teal), talonavicular (navicular; blue), and subtalar (calcaneus; purple) joints for all participants (n_Participants_ = 9; n_Trials_walk/run_ = 1; n_Trials_hopping_ = 3). Participants’ orientation values for some bones might be missing if the bone could not be tracked or was ≤ 5°. In these instances, there are no data presented in the graphs (indicated with an X).

**Figure 5:**
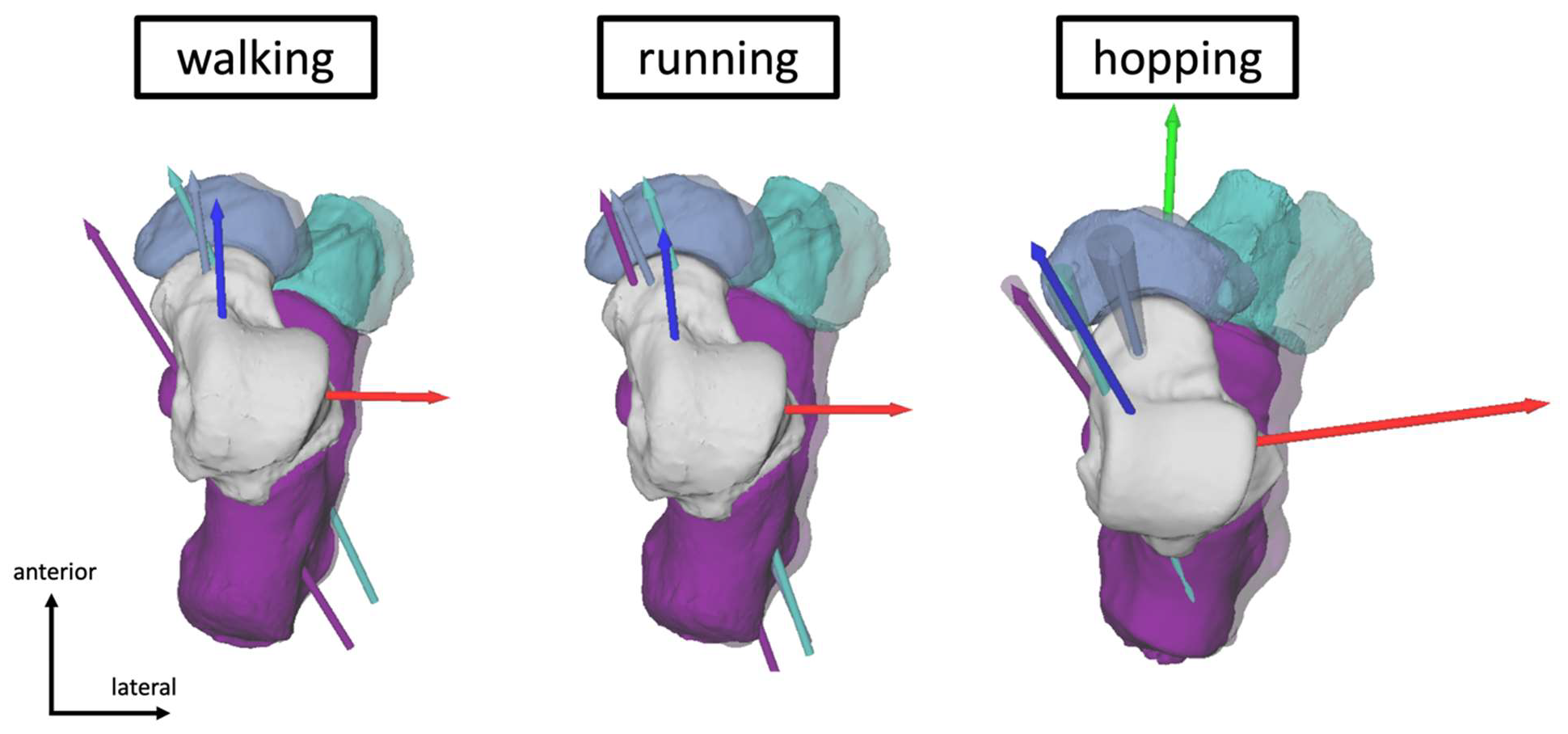
Right foot; superior view. Exemplary rotation axes during push-off across different modes of locomotion (columns) for the talonavicular (navicular; blue), talocuboid (cuboid; teal), and subtalar (calcaneus; purple) joints for two participants (note that the participant for walking and running is not the same as for hopping). The cone around the rotation axis during hopping reflects the standard deviation across three trials. The talus coordinate system is reflected in red, green, and blue (x-, y-, z-axis respectively). Transparent bones indicate the posture of the bones during the transition point while solid bones indicate the posture of the bones during toe-off. If the cuboid was missing, the data could either not be tracked or the total amount of rotation was ≤5° and therefore excluded.

Across locomotor modes, the calcaneus consistently rotated substantially less than the cuboid or navicular (Table 1). These results were significant for hopping (calcaneus rotates significantly less than the cuboid and navicular; p ≤ 0.0078, Supplement Table S3) and running (calcaneus rotates significantly less than the navicular; p = 0.0039, Supplement Table S2). The navicular and cuboid showed no significant differences in their rotation magnitudes (Figure 5 & Table 1), despite the cuboid consistently rotating on average about 2-3° (range -9 – 1°) less than the navicular across locomotor modes and while rotating about similarly oriented rotation axes.

We also found that most of each joint’s motion occurred about the morphological subtalar axis (Figure 4 & 5), with average orientation magnitudes ranging from 0.86 to 0.95 (Table 1). In other words, approximately 90% of the total rotation of the talonavicular, talocuboid, and subtalar motion occurs about the morphological subtalar axis.

## 4. Discussion

The objective of our study was to determine if the midtarsal complex and subtalar joint behave similarly during locomotion and if this motion is consistent with talus articular morphology. To accomplish our objectives, we used highly accurate BVR equipment to measure bone morphology in motion during walking, running, and hopping. We found that the subtalar, talonavicular, and talocuboid joint axes are closely aligned with each other and with the morphological subtalar axis regarding their orientations across both gait phases. Although the range of motion and the location of the axes differed among bones (Figure S8), these findings suggest that the midtarsal and subtalar joints behave as a cohesive bone complex that is driven by talus articular surface morphology. This behavior cannot be seen by analyzing all three joints independently and with individual coordinate systems. Our findings are not surprising for the calcaneus and navicular, which directly articulate with the talus. However, it is noteworthy that the cuboid is separated from the talus by the calcaneus, yet still primarily rotates relative to the talus about an axis oriented similarly to the morphological subtalar axis. This suggests that using the talus as a common reference bone for midtarsal and subtalar complex motion might be a good approach to investigate non-pathological but also pathological locomotion patterns in the future.

All bones substantially rotated about an axis aligned with the morphological subtalar axis during walking, running, and hopping. Interestingly, despite sharing a commonly oriented axis of motion, the three bones exhibited differing magnitudes of rotation and some variation in the location of the rotation axis locations, reflecting high levels of inter-subject variation. The large inter-subject variation in axis location (Figure S8) did not allow us to detect a pattern. It is unclear whether the observed variation is due to morphological variability or measurement errors; therefore, we did not make inferences on axis location. Our findings are consistent with previous bone-pin and BVR studies showing that the navicular and cuboid rotate similarly [7,15,16,25], with the caveat that the navicular rotates on average 12% more than the cuboid during the push-off phase. Additionally, we observed that the navicular and cuboid bones display substantially greater magnitudes of rotation than the calcaneus during push-off (∼35-60% more depending on locomotion type), which is similar to previous literature [15,17,25]. These data contrast with Elftman and Manter’s [21] suggestion that the midtarsal joints lock (i.e. does not move) when the talonavicular and calcaneocuboid joint axes converge from midstance to push-off, converting the foot into a “rigid” or stiff lever for push-off. Others have also demonstrated that the midtarsal and subtalar joints do not lock but rather display great mobility [14–16,25]. Our results further strengthen the evidence against midtarsal locking during push-off. Consequently, we suggest embracing the notion that the human foot is highly mobile throughout locomotion, facilitated by a kinematic axis aligned with the morphological subtalar axis, and does not behave as a single rigid unit.

We are intrigued by our findings that the navicular, cuboid, and calcaneus rotate different amounts about a commonly oriented axis that is defined by subtalar and talonavicular joint facet morphology [23]. Since the talocuboid and subtalar joint motions are less than talonavicular motion (particularly the subtalar joint), we speculate that these joints may be constraining the navicular’s motion path relative to the talus. Since the talonavicular has been described as an elliptical paraboloid articulation [39], it is reasonable to assume it possesses more potential degrees of freedom than the saddle-shaped articular surfaces of the calcaneocuboid and subtalar joints [21]. We believe this system may be analogous to the engineering design of a dual track slider, where one groove has a tight tolerance to define the motion (subtalar joint and to a lesser degree the calcaneocuboid joint) and the second groove has a looser tolerance and is deemed the follower (talonavicular joint). This configuration is advantageous because it prevents the entire dual track slider from locking. Hence, the navicular might be adjusting and potentially compensating for limited motion of the other joints as necessary. That said, this interpretation remains speculative and requires further exploration, as we cannot determine cause and effect with the current methods.

The morphology of a bone’s articular surface a key determinant of how that bone or joint moves during locomotion; however, osseus morphology is rarely considered in dynamic evaluation of human foot function [29,40,41]. Recent statistical shape modelling (morphometrics) studies of the human midtarsal bones have highlighted the marked inter-subject variability in the shape of the articular surfaces of the talus [11,13]. Our morphological axis approach captures important shape features and its variation among individuals. It is possible that variations in shape (and joint axis orientations and locations) may explain the wide inter-subject variation in the magnitude of rotation, with particular shape features enabling greater, or restricted rotation about a commonly oriented axis. We did not control for walking and running speeds in our study, so it is also possible that variation in gait speed may also contribute to varying magnitudes of rotation. However, we controlled for frequency in hopping, and the intra-individual consistency in rotation magnitude between walking, running, and hopping suggests speed effects may be minimal. For example, individuals who exhibited lower rotation magnitudes in walking typically also reported lower rotation magnitudes in running. Understanding the contributing factors to inter-subject variability is important for clinicians and anthropologists when determining boundaries for healthy versus pathological midtarsal complex and subtalar joint function, planning clinical treatments, or when comparing midtarsal kinematics across species.

### 4.1. Limitations

The acquisition and processing of data using BVR is time-intensive, which is why our sample size is relatively small. Due to the limited field of view and the stationary setup of the BVR, it is not possible to repeatedly capture the entire stance phase for all foot bones during walking and running, resulting in missing data. However, we were able to capture more than 90% of stance on average (Supplement Figure S1), which should suffice for most biomechanical and clinical applications. As we studied sagittal plane dominant tasks, it is unclear how our results extend to tasks with substantial out of plane ankle motion, such as cutting. Moreover, bone occlusion (bony overlap in the capture volume) can make bone tracking challenging or impossible, particularly in the midfoot region. These limitations make it difficult to provide a complete dataset of cuboid kinematics for each mode of locomotion and participant. Therefore, researchers may need to prioritize which bones are imaged to address their research question and optimize their BVR setup accordingly. Finally, it is worth noting that we collected the walking and running data while participants wore minimalist shoes to minimize discomfort and promote their regular gait pattern, assuming that participants were not habitual barefoot runners. Considering that the shoe displayed minimal cushioning, we believe it had little impact on bone motion compared to the hopping data set, which was performed barefoot.

### 4.2. Future Application

Based on the findings presented in this study, we speculate that morphological variations in subtalar and talonavicular joint articular surfaces may have substantial effects on the alignment of the morphological subtalar axis and, thus, the kinematics of the entire midtarsal complex and subtalar joint. Changes in osseous structure occurring due to conditions such as osteoarthritis may lead to a progressive sequence of structural and functional impairments. At the opposite end of the lifespan, it is also possible that developmental plasticity associated with childhood (and lifetime) physical activity may also play a considerable role in how our feet function through adulthood into our elderly years. This uniaxial representation of the midtarsal complex and subtalar joint might simplify future implant design in subtalar joint replacements or talus arthroplasties. Moreover, looking at midtarsal complex and subtalar joint kinematics relative to the talus might be an approach to distinguish healthy from pathological midtarsal complex and subtalar joint function instead of interpreting joint motion independently (e.g., calcaneocuboid joint). This would streamline the interpretation of midtarsal complex and subtalar joint kinematics and clearly identify individuals with deviating rotation axes and movement patterns.

Our findings might also be of interest for biological anthropologists who suggest that morphological variations in foot bones, particularly the talus, played a key role in the evolution of foot function, transitioning from non-bipedal to bipedal locomotion [42–45]. Human talus morphology exhibits remarkable differences in size and shape compared to chimpanzees (our closest living relatives) and early hominins [43–45]. We expect that the changes of the talonavicular and subtalar facets specifically may be critically important morphological features as they determine the orientation and location of the morphological subtalar joint axis. Studies [21,42,44,46] show modern humans have increased talar head torsion, a lower talar neck angle, and less curved (flatter) talonavicular and subtalar articular facets compared to chimpanzees and early hominins. A subtalar and talonavicular surface with less curvature may result in a steeper (more superiorly oriented) morphological subtalar axis. This, in turn, may be responsible for the limited inversion/eversion motion of the foot relative to the lower leg in modern humans. It is possible that the morphological changes were advantageous for example by providing increased stability during bipedal locomotion. Understanding the link between midtarsal and subtalar joint morphology and function is crucial, especially when fossil features are difficult to link directly with dynamic motion.

## 5. Summary

We have shown that the calcaneus, navicular, and cuboid bones move about a commonly oriented axis that coincides with the morphological subtalar axis during walking, running, and hopping. Furthermore, we observed substantial midtarsal complex and subtalar joint movement during landing and push-off in different modes of locomotion, suggesting a considerable degree of mobility throughout the stance phase. Lastly, our results underscore the importance of understanding both bone motion and morphology and the potential implications for developing targeted foot treatments for pathological gait characteristics and providing further details about the evolution of bipedalism.

