## Supplemental Data for "Human *in vivo* midtarsal and subtalar joint kinematics during walking, running, and hopping"

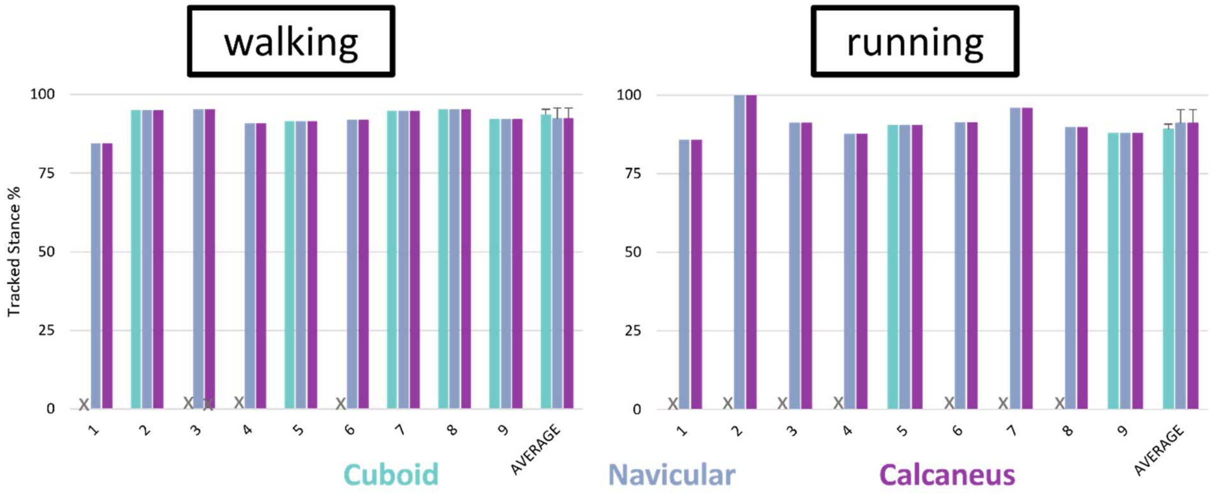


*Figure S1: Tracked Stance in % for all three bones of the tarsal complex relative to the talus bone. If data is missing, the bone was not tracked for the trial due to osseous occlusion or because the total rotation was ≤ 5°.*

*Table S1: One-way ANOVAs for walking to compare whether there are significant differences between the three bones (navicular, cuboid, and calcaneus) during landing and push-off phases the orientation in the morphological subtalar axis direction and total rotation. Post hoc tests are included if the group comparison was significantly different (indicated with an asterisk).*

| ***WALKING*** | | *p-value* | *alpha* |
|---|---|---|---|
| *Landing* | Total rotation | 0.1250 | 0.025 |
| | Orientation | 0.8066 | 0.025 |
| *Push-off* | Total rotation | 0.1041 | 0.025 |
| | Orientation | 0.3207 | 0.025 |

*Table S2: One-way ANOVAs for running to compare whether there are significant differences between the three bones (navicular, cuboid, and calcaneus) during landing and push-off phases the orientation in the morphological subtalar axis direction and total rotation. Post hoc tests are included if the group comparison was significantly different (indicated with an asterisk).*

| ***RUNNING*** | | *p-value* | *alpha* |
|---|---|---|---|
| *Landing* | Total rotation | 0.07 | 0.025 |
| | Orientation | 0.99 | 0.025 |
| *Push-off* | Total rotation | 0.005*<br><br>Post-hoc<br>cal vs cub: 0.5<br>cal vs nav: 0.004*<br>cub vs nav: 1 | 0.025<br><br>Post-hoc<br>0.02 |
| | Orientation | 0.78 | 0.025 |

*Table S3: One-way ANOVAs for hopping to compare whether there are significant differences between the three bones (navicular, cuboid calcaneus) during landing and push-off phases the orientation in the morphological subtalar axis direction and total rotation. Post hoc tests are included if the group comparison was significantly different (indicated with an asterisk).*

| ***HOPPING*** | | *p-value* | *alpha* |
|---|---|---|---|
| *Landing* | Total rotation | 0.0026*<br><br>Post-hoc<br>cal vs cub: 0.004*<br>cal vs nav: 0.004*<br>cub vs nav: 0.6 | 0.025<br><br>Post-hoc<br>0.02 |
| | Orientation | 0.3 | 0.025 |
| *Push-off* | Total rotation | 0.01*<br><br>Post-hoc<br>cal vs cub: 0.008*<br>cal vs nav: 0.008*<br>cub vs nav: 0.04 | 0.025<br><br>Post-hoc<br>0.0167 |
| | Orientation | 0.2 | 0.025 |

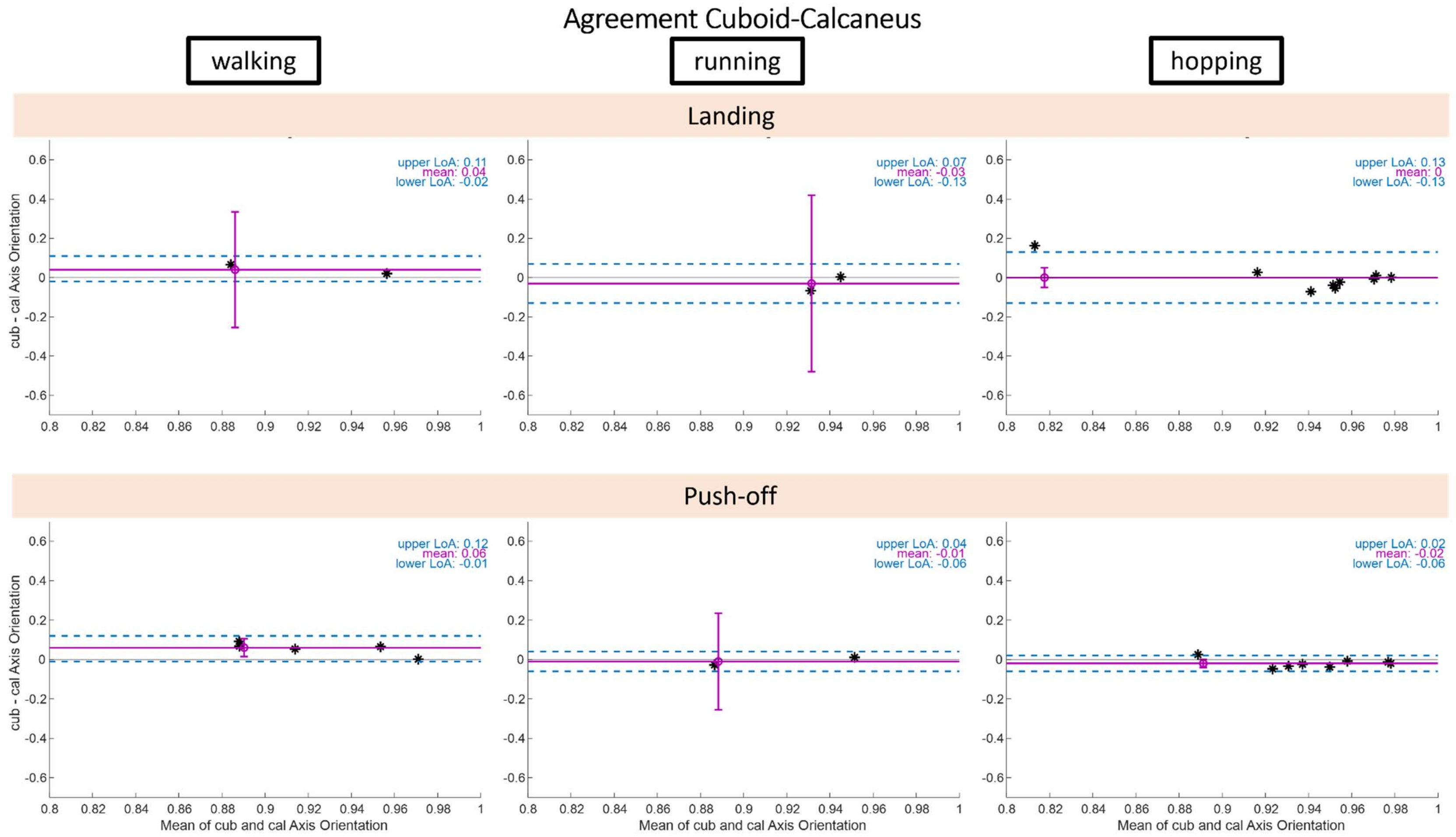


*Figure S2: Bland Altman plots for comparison of orientation between cuboid and calcaneus bone for all three modes of locomotion (columns). No differences between both bone orientations were observed except during the push-off phase in walking (calcaneus axis is oriented less in the subtalar joint axis orientation than the cuboid). The limits of agreement (LoA in blue) indicate good agreement and the error bar indicates the 95% confidence interval of the mean (purple).*

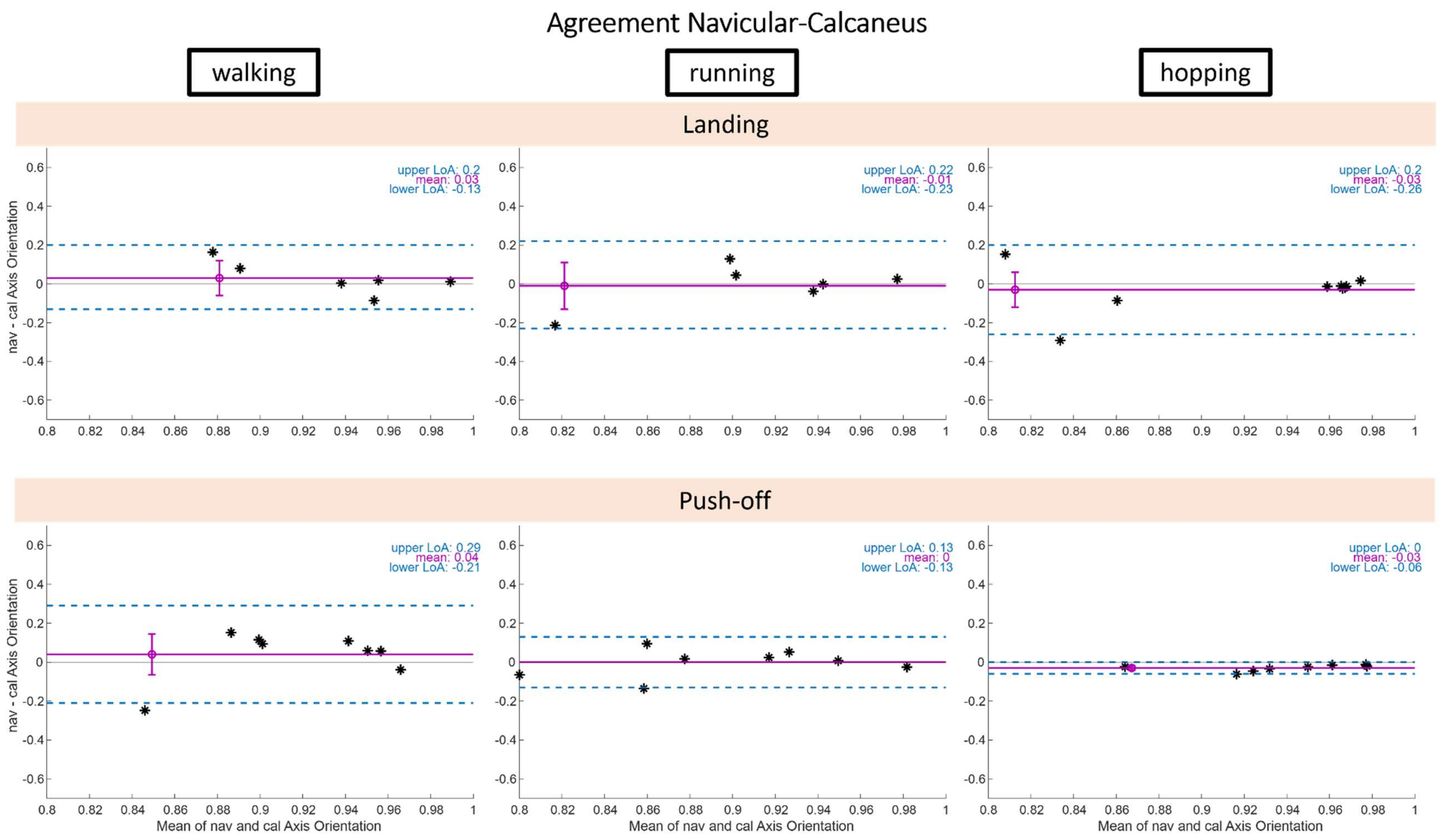


*Figure S3: Bland Altman plots for comparison of orientation between navicular and calcaneus bone for all three modes of locomotion (columns). No differences between both bone orientations were observed except during the push-off phase in hopping (navicular axis is oriented less in the subtalar joint axis orientation than the calcaneus). The limits of agreement (LoA in blue) indicate good agreement and the error bar indicates the 95% confidence interval of the mean (purple).*

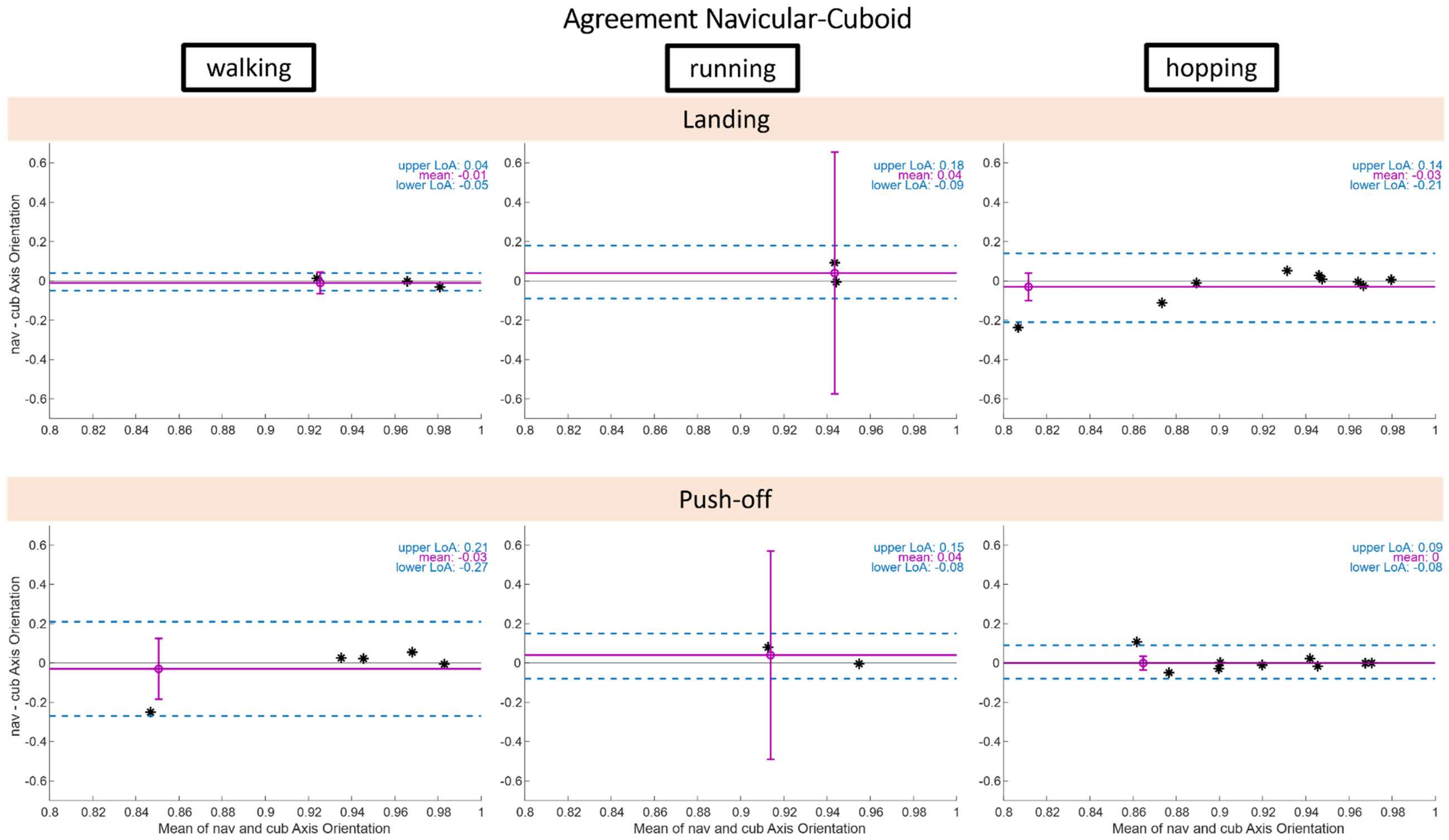


*Figure S4: Bland Altman plots for comparison of orientation between navicular and cuboid bone for all three modes of locomotion (columns). No differences between both bone orientations were observed. The limits of agreement (LoA in blue) indicate good agreement and the error bar indicates the 95% confidence interval of the mean (purple).*

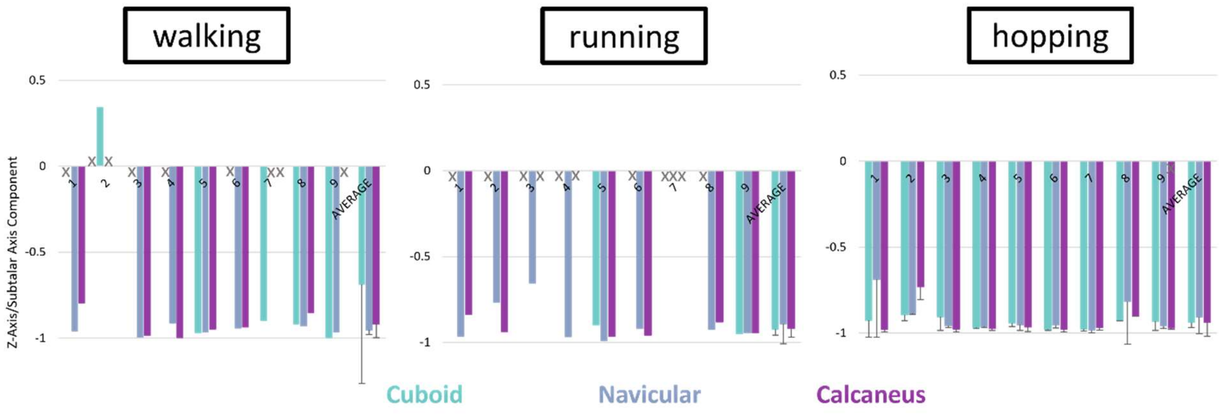


*Figure S5: Total rotation in degree during landing phase of different modes of locomotion for the talocuboid (cuboid; teal), talonavicular (navicular; blue), and subtalar (calcaneus; purple) joints for all participants ($n_{Participants}$ = 9; $n_{Trials\_walk/run}$ = 1; $n_{Trials\_hopping}$ = 3). Participants' total rotation for some bones might be missing if the bone could not be tracked or was ≤ 5° in these instances there is no data presented in the graphs (indicated with an X).*

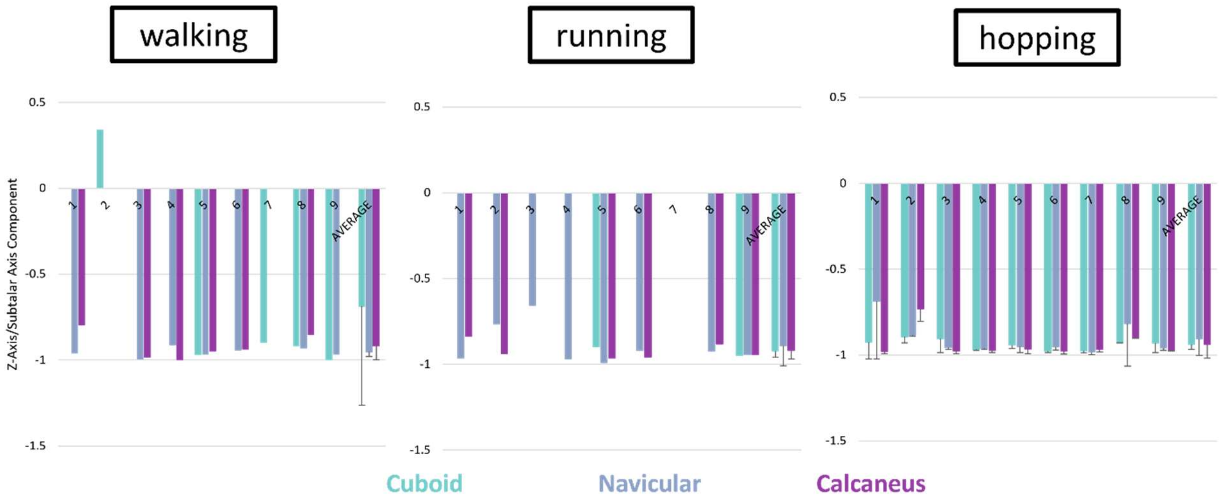


*Figure S6: Orientation of helical axis of motion in the subtalar axis direction (z-axis) during landing phase of different modes of locomotion for the talocuboid (cuboid; teal), talonavicular (navicular; blue), and subtalar (calcaneus; purple) joints for all participants ($n_{Participants}$ = 9; $n_{Trials\_walk/run}$ = 1; $n_{Trials\_hopping}$ = 3). Participants' orientation values for some bones might be missing if the bone could not be tracked or was ≤ 5° in these instances there is no data presented in the graphs (indicated with an X).*

*Table S4: Spatial angle (median ± standard deviation) between the subtalar axis (z-component) and participants per bone across all three locomotion tasks.*

| | ***WALKING*** | | ***RUNNING*** | | ***HOPPING*** | |
|---|---|---|---|---|---|---|
| | *Landing* | *Push-off* | *Landing* | *Push-off* | *Landing* | *Push-off* |
| *Calcaneus* | 19.7 ± 12.3 | 25.2 ± 9.2 | 21.0 ± 7.0 | 25.8 ± 13.2 | 13.0 ± 10.1 | 16.6 ± 6.3 |
| *Navicular* | 16.3 ± 6.1 | 16.9 ± 13.6 | 21.1 ± 14.4 | 24.9 ± 12.8 | 15.9 ± 10.0 | 20.2 ± 5.9 |
| *Cuboid* | 23.5 ± 42.2 | 19.9 ± 5.5 | 22.5 ± 5.2 | 23.1 ± 8.7 | 18.7 ± 5.5 | 19.2 ± 6.6 |

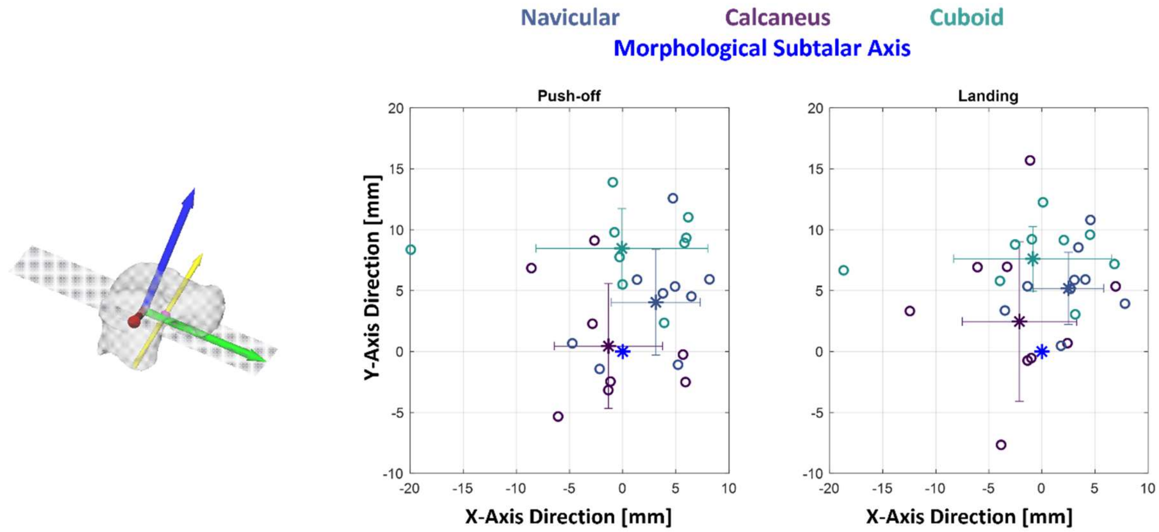


*Figure S7: This figure serves as an example of the spread in rotation axis location across participants and bones. On the right side, an example of a rotation axis (yellow arrow) and the intersection point (pink) of the axis with the xy-plane is illustrated for better understanding. The graphs show the intersection points of the rotation axes during hopping (push-off phase on the right, landing phase on the left) with the xy-plane. Each participant's average across trials is reflected by a circle and the average intersection location across each bone is indicated with an asterisk and the standard deviation is reflected as error bars in x- and y-direction. The subtalar axis is indicated as a blue asterisk at the origin. A similar spread was observed during walking and running tasks.*
